# Programmable acoustic single cell manipulation with model-free machine learning

**DOI:** 10.64898/2026.06.29.735220

**Authors:** Alexander Edthofer, Guilherme J. Perticarari, Selma Helvelius Bounja, Thierry Baasch

## Abstract

Precise, non-invasive manipulation of individual living cells remains a central challenge in biomedical science, with far-reaching implications for single-cell analysis, tissue engineering, and the study of cell–cell interactions. Here, we report the first demonstration of single-cell control using bulk acoustic standing-wave acoustofluidics with closed-loop feedback. We introduce **VeLO** (**Ve**ctor-based **L**ocal **O**ptimization), a model-free, reinforcement learning–inspired algorithm that enables programmable two-dimensional manipulation of individual cells using a single piezoelectric transducer. Without prior calibration or physical modeling, VeLO learns system dynamics online from acoustically induced cell displacements and automatically adapts to nonlinear, time-varying conditions. We achieve robust control across multiple cell types (DU-145, Jurkat, K-562) and independent manipulation of multiple cells, including controlled cell–cell contact. By combining simplicity of hardware with autonomous, adaptive control, this approach establishes multimodal acoustofluidics as a versatile tool for label-free, high-precision single-cell manipulation.

## 1. Introduction

Manipulating individual cells with high spatial and temporal precision is a long-standing objective in biomedical research and a foundational capability for modern quantitative biology and medicine. Deterministic single-cell control is the basis of a wide range of applications, including single-cell omics and cell selection for downstream analysis,^[1,2]^ the bottom-up assembly of tissues and organoids,^[3]^ and the controlled interrogation of cell-cell interactions and cellular mechanics in immunology, cancer biology, and developmental systems.^[4]^ The recent rise in single-cell genomics has revealed the enormous functional heterogeneity of seemingly identical cells,^[1,2]^ making it clear that technologies capable of positioning, isolating, and perturbating individual cells with high fidelity are central to enabling next generation experimental biology.

Over the last four decades, micro-and nanoscale manipulation has been shaped by the development of contact-free force fields. Among these, optical tweezers have enabled the trapping and manipulation of microscopic objects with nanometer precision ^[5]^ and have become indispensable for probing molecular motors, cell mechanics, and biophysical interactions,^[6]^ resulting in the 2018 Nobel Prize in Physics.

While various techniques exist for single-cell manipulation, each comes with fundamental trade-offs. Micropipettes and microgrippers offer direct mechanical control but require physical contact, which can compromise viability, perturb cellular state, and limit throughput and automation. ^[7–9]^ Dielectrophoresis offers non-label and non-contact control but requires specific medium requirements.^[10]^ Optical tweezers provide non-contact manipulation with submicron precision but can induce photothermal or photochemical damage,^[11]^ require complex and costly optical setups, and often provide limited trapping forces for large, weakly refracting, or highly motile cells, increasing the likelihood of escape.^[3]^ As a result, achieving reliable, gentle, and scalable manipulation of individual live cells remains a central unsolved challenge.

Acoustic manipulation, in particular acoustic tweezers, has emerged over the past decade as a promising alternative.^[12–14]^ By exploiting sound-induced radiation forces and streaming flows,^[15,16]^ acoustic fields can manipulate particles and cells in a label-free, contact-free, and highly biocompatible manner at power densities orders of magnitude lower than optical methods. Acoustic approaches are inherently scalable and compatible with microfluidic integration, leading to rapid progress in acoustofluidics for trapping, patterning, sorting, and manipulating cells.^[17–21]^ More recently, a new generation of acoustic tweezers has demonstrated increasingly sophisticated capabilities, including complex boundary-free trapping landscapes,^[22]^ size-insensitive single-cell manipulation and mechanical phenotyping,^[23]^ high-throughput single-cell analysis,^[13]^ independent multi-particle control and cell deformation,^[14,24]^ resonant-field engineering,^[25]^ and streaming-based transport and assembly.^[26]^ Acoustic tweezers are now widely seen as a mature and rapidly expanding toolbox for precision biology and medicine.^[27]^

Most existing acoustic manipulation approaches rely on static or quasi-static pressure fields generated by carefully designed transducer geometries or mechanically reconfigurable platforms.^[12,26]^ These approaches fundamentally couple device geometry to functionality and often limit flexibility, reconfigurability, and independent control. While they repeatedly have shown their many uses,^[12–14, 22–27]^ many tweezer-based strategies are constrained to one object per trap or require complex hardware scaling to achieve parallelization, making selective manipulation within dense and heterogeneous cell populations challenging.

The combination of acoustic actuation and feedback control has been utilized for trapping, navigation, and steering of microrobots in vivo, including in demanding environments such as the mouse brain vasculature^[28]^ and autonomous microrobotic systems capable of adaptive, goal-directed behavior,^[29–31]^ showing clear applications for targeted intervention and precision therapy.^[32]^

Multimodal acoustofluidics offers a promising route to overcome these limitations. Based on bulk acoustic wave (BAW) resonators, multimodal acoustofluidics exploit that a single microfluidic cavity supports many acoustic resonance modes.^[33]^ By dynamically switching or mixing these modes, the acoustic pressure landscape can be reshaped in time, enabling object-specific control within a single, fixed device.

Early work has shown that machine-learning-based, model-free control can harness this high-dimensional actuation space to transport and sort particles in complex and adaptive ways. ^[34, 35]^ However, a central challenge remains the reliable and precise control of living cells. In practice, the acoustic field inside a device is difficult to predict and maintain due to fabrication tolerances, transducer coupling, and environmental variations such as temperature-dependent changes in sound speed and fluid viscosity.^[16]^ These effects make purely model-based approaches unreliable and motivate adaptive, feedback-driven strategies.

In this article, we present two key advances. First, we demonstrate, for the first time, deterministic manipulation of individual live cells using bulk acoustic wave multimodal acoustofluidics. We show precise two-dimensional control of single cells from multiple cell lines, as well as independent, simultaneous control of multiple cells within the same device, including bringing two cells into direct contact. In contrast to multitransducer or phased-array approaches, this strategy relies on a single transducer, reducing hardware complexity while retaining a high degree of control. Secondly, we introduce a new reinforcement learning-inspired control algorithm, termed VeLO (Vector-based Local Optimization). VeLO preserves the model-free adaptability of ε-greedy previously applied for the control of polystyrene particles and droplets^[35–37]^ while incorporating a velocity vector-based mode-mixing approach,^[38]^ resulting in improved control speed, robustness, and accuracy.

Together, these results establish multimodal acoustofluidic manipulation as a robust and accessible platform for single-cell control. By combining a simple microfluidic architecture with adaptive, feedback-driven control, this approach brings manipulation conceptually closer to what optical tweezers have represented for the past four decades, while offering a complementary regime of forces, scalability, and biocompatibility. More broadly, this work expands on the combination of acoustics, robotics, and machine learning and contributes to ultrasound-driven micromanipulation becoming a routine component of lab-on-a-chip systems for both fundamental biology and translational biomedical applications.^[29–33, 35–38]^

## 2. Results

### 2.1. Principle and platform

Acoustic pressure fields arise from sound-induced vibrations of a material or medium. As with other physical fields, acoustic pressure fields exhibit spatial minima and maxima whose locations are strongly frequency dependent and become most pronounced when the actuation frequency matches the resonance modes of the system. A classic macroscopic illustration of vibrating systems inducing particle motion are Chladni plates, in which particles such as sand redistribute into characteristic nodal patterns upon resonant excitation of a vibrating plate.^[39]^

The same physical principles apply at the microscale. In microfluidic devices, acoustic pressure fields can be generated within fluid-filled cavities by actuating an attached piezoelectric transducer.^[18–21]^ Suspended objects, including biological cells, experience forces arising from gradients in the acoustic field and migrate toward stable equilibrium positions. For particles that are small compared to the acoustic wavelength, the time-averaged acoustic radiation force can be derived from the Gorkov potential *U*:^[15]^

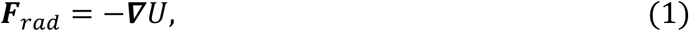

where the potential is given by

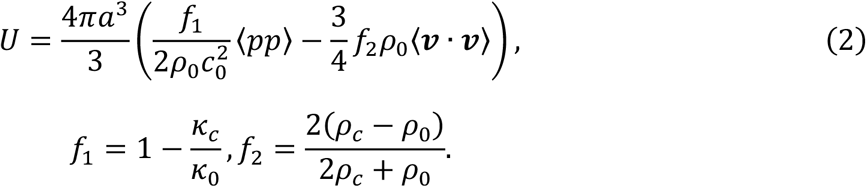

Angle brackets denote time averaging, *a* is the particle radius, *p* the acoustic pressure, *v* acoustic velocity, *c_0_* the speed of sound in the fluid. The monopole *ƒ_1_* and dipole *ƒ_2_* scattering coefficients are material properties and are a function of the cell (*c*) and surrounding fluid (0) density (*ρ*) and compressibility (*κ*).

It is important to note that living cells differ fundamentally from commonly used model particles such as polystyrene beads, exhibiting lower scattering and are more heterogenous, making them more difficult to manipulate.

We implement a feedback control algorithm that selects appropriate actuation modes at each iteration. A schematic of the control loop is shown in **Figure 1**A. Prior to closed-loop operation, the experiment is initialized by selecting target cells via a custom MATLAB graphical user interface (GUI) with a live microscopy feed. The same interface is used to define target positions and to configure experimental parameters, including the actuation time step *Δt*, input voltage, and the set of accessible actuation frequencies. The control algorithm works as follows:

**Figure 1.**
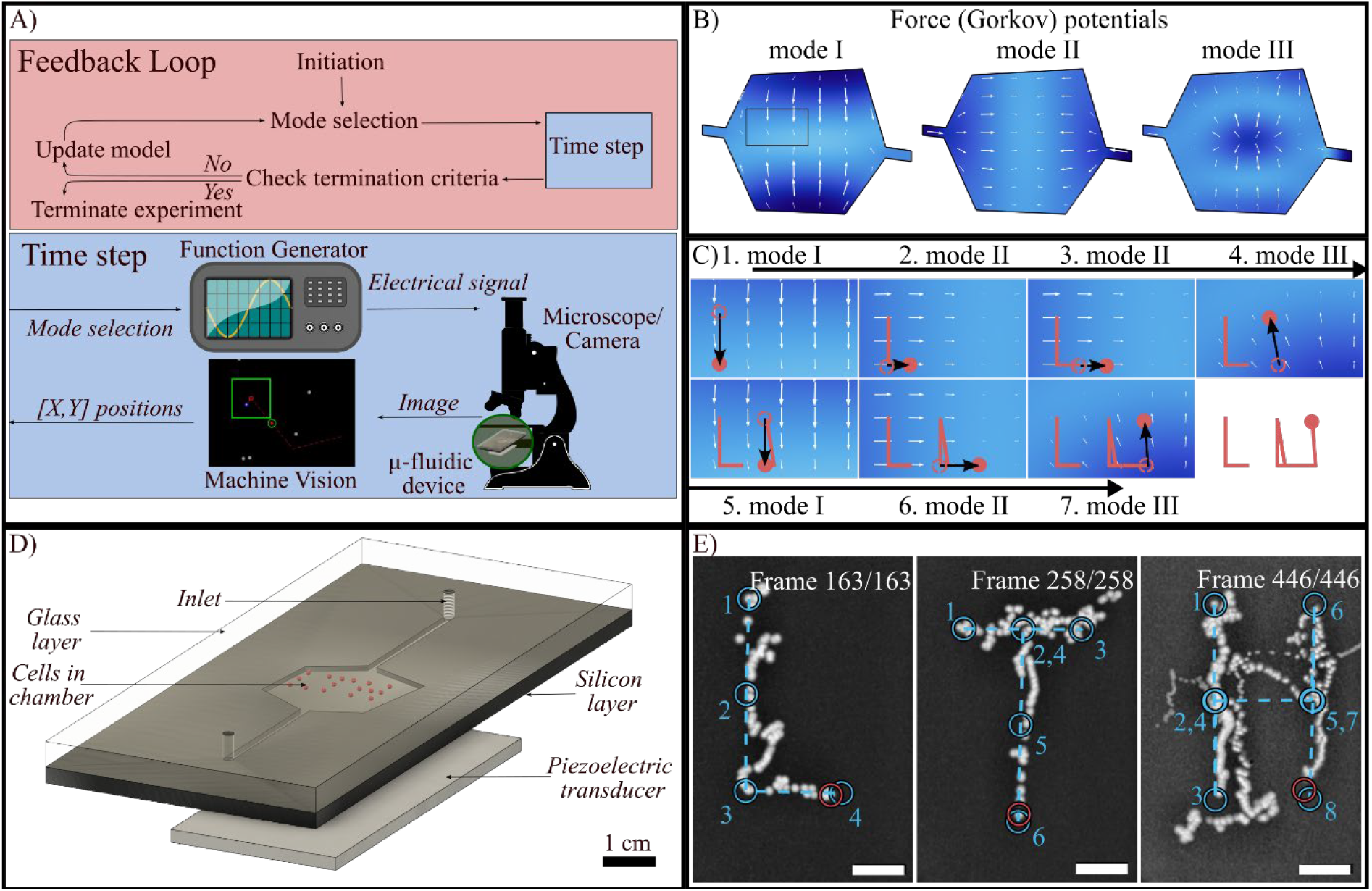
Algorithm logic and platform. (A) Flowchart over the feedback loop of the control system (top), and how it couples to the timestep taken in the physical platform consisting of a function generator, the microfluidic device, and a microscope with camera (bottom). (B) COMSOL simulations of the Gorkov potential inside of the microfluidic when actuating the chamber at 0.20 (mode I), 0.18 (mode II), and 0.40 (mode III) MHz. Regions of high and low potential are indicated by dark and light blue, respectively. White arrows show the negative gradient of the Gorkov potential. Black box is the region displayed in C. (C) A series showing the movement of a cell (brown dot, displacement indicated by black arrow), due to the acoustic radiation force. By alternating between modes, the cell is moved to spell out “LU”. (D) The microfluidic device. (E) An individual DU-145 prostate cancer cell is controlled to write “LTH”. The image was created by merging all experimental frames. Numbered blue circles are the intermediate target positions, red circles the final cell position. Dashed blue lines are the shortest path. Scale bar is 100 µm.

#### Initialization

For a system with modes *m*∈{1,…,*M*}, cells *c* ∈ {1, …, *C*} with current positions 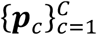 and target positions 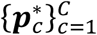, and hyperparameters *γ* ∈ (0,1), and *λ* > 0, the displacement model *W*_*m,c*_ ∈ ℝ^*D*=2^ is initiated by actuating each mode once for a time step *Δt* and track the cell displacement. The directional estimate *u*_*m,c*_ and the uncertainty model *V*_*m,c*_ are initiated as empty vectors.

#### Main Loop

The control loop consists of the following steps, and repeats until all cells have reached their final target:

1. *Compute target displacements*

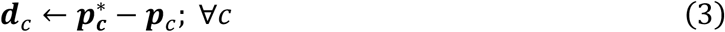
2. *Compute optimal mode mixture*
Let

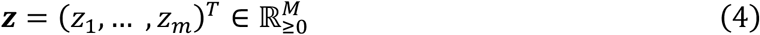

Denote the non-negative mode mixture weights. Compute the minimized mixture

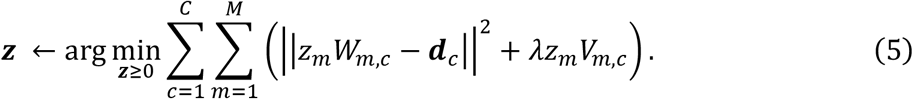
3. *System actuation*
  - Sample 5 modes independently according to the categorical distribution of **z**.
  - Apply the selected modes sequentially for a duration of *Δ*.
4. *State observation*
  - Track cell positions via machine vision
5. *Model and target state update* For each mode m and cell c:
  1. Calculate observed displacement:

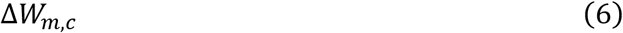
  2. Update model:

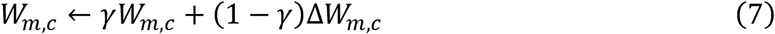
  3. Update directional estimate:

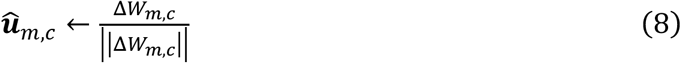
  4. Update model uncertainty:

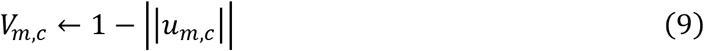
  5. Update target position 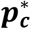 if target was reached.

Thus, the control algorithm operates without the need for any prior knowledge of the system dynamics and instead learns online based on the cell’s displacements after actuation.

Figure 1B shows finite-element simulations (COMSOL Multiphysics) of the Gorkov potential within a model microfluidic chamber at different actuation frequencies, further explained in Supplement A. Cells migrate along the negative potential gradient, illustrated by the white arrows. Figure 1C demonstrates how alternating between these frequency-dependent potential landscapes enables controlled cell trajectories, exemplified by writing the letters “LU”. All experiments were conducted in a microfluidic device with irregular chamber geometry. This design suppresses spatial symmetry in the acoustic pressure fields, which otherwise may introduce barriers to cell motion. For the same reason, the single piezoelectric transducer was mounted off-center near a corner of the device. A schematic of the device is shown in Figure 1D.

In Figure 1E, we demonstrate closed-loop control of a single live DU-145 prostate cancer cell suspended in cell buffer, guided to trace the initials of Lunds Tekniska Högskola, LTH. The total target path for all letters combined was 2.58 mm, and required 867 control iterations, with each iteration containing a control input of 5 frequencies played sequentially. The experiment was completed in less than 2.5 hours. Notably, the cumulative actuation time was only 22 minutes, corresponding to 4335 actuation steps across 867 control iterations, highlighting the substantial potential for further optimization and acceleration of this approach. A video of the experiment can be found in Supplement B. Any experimental parameters in the manuscript fall between 0.3-0.7 s for *Δt*, 10-23 V for *V*_in_, *γ* = 0.2, *λ* = 0.0005 and a set of 30-40 discrete actuation frequencies uniformly distributed between 1.3 – 3.2 MHz. The shifts in experimental settings were mainly due to differences in energy transfer efficiency between the piezo and the microfluidic device caused by replacing and regluing the piezo. A table with the experimental values for each individual experiment can be found in Supplement C.

### 2.2. Algorithm characterization

Two key performance metrics of any closed-loop manipulation platform are the speed at which a prescribed task can be completed and the accuracy with which target positions can be reached. Rather than relating experimental outcomes to input parameters such as voltage amplitude, acoustic power, or actuation duration *Δt*, the mobility is directly characterized through the average displacement per control step. This metric captures the intrinsic dynamics of the coupled system, is directly observable experimentally, and enables comparison across devices with different geometries or actuation characteristics.

We therefore quantify algorithm performance by evaluating the number of control steps required to traverse a normalized distance of 1 mm as a function of the average step size. To quantify the accuracy of the algorithm, the bounding area between the cell trajectory and the direct path between target positions is measured. Further on in this work, this bounding area will be referred to as the error area *A*_*error*_.

This analysis was performed numerically using a simplified simulation framework, described in more detail in Supplement D. The simulated microfluidic chamber was assumed to be rectangular, and the accessible acoustic modes consisted of combinations of first to sixth order standing waves in both the x- and y-directions, yielding 36 distinct pressure field configurations. Aside from these simplifications, the physical parameters were chosen to match those of non-adherent DU-145 prostate cancer cells suspended in phosphate-buffered saline, with a = 7.5 µm,*ƒ_1_* = 0.12 and *ƒ_2_*= 0.038. The values were taken from literature.^[40]^

In each simulation, the control algorithm guided a single cell sequentially between three predefined target positions before returning it to its starting location, creating a triangle. The acoustic pressure amplitude was incrementally increased between simulations, thereby systematically increasing the average displacement per step while maintaining control logic. To show general validity of the numerical results, additional simulations using a target trajectory which was rotated 60° degrees were performed.

Figure 2 summarizes the simulation results, showing the number of steps *A*_*error*_ as function of the average step size. The number of steps decreases sharply with increasing step size, reflecting more efficient cell manipulation, until a minimum is reached at an average displacement of approximately 3 µm. This value corresponds to roughly one-fifth of the simulated cell diameter (15 µm). Beyond this point, further increases in step size led to a rise in the number of required steps, as the algorithm increasingly overshoots the target region and struggles to converge within the prescribed positional threshold. In parallel, *A*_*error*_ increases with the average step size, indicating a trade-off between speed and trajectory fidelity.

**Figure 2.**
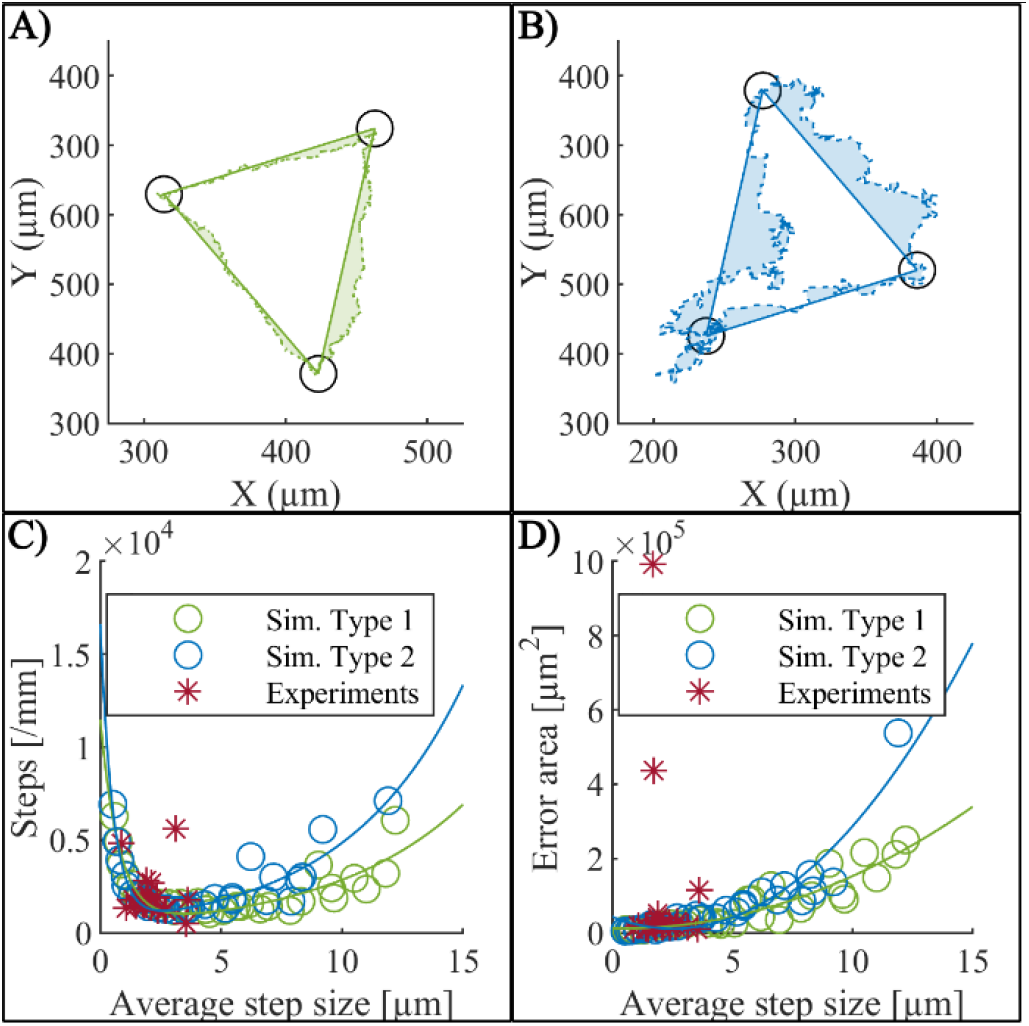
Numerical characterization of cell control. (**A**) Tracing Sim. Type 1, with an average step size of 0.85 µm, requiring 3117 steps. (**B**) Tracing Sim. Type 2 with an average step size of 4.32 µm, requiring 1517 steps. Dotted lines are the trajectory, shaded areas *A*_*error*_. (**C**) The number of steps needed to traverse 1 mm of a trajectory as a function of the average step size. (**D**) A_error_ between the cell path and the direct line between start and target as a function of the average step size. Blue and green rings are numerical results obtained from Sim. Type 1 and Sim. Type 2, respectively; lines are quadratically fitted functions. Red stars are data extracted from experiments found in Supplement C. The data indicates an optimal operational region for an average step size between 2 and 5 µm, which minimizes the number of steps needed to complete a control task with high trajectory accuracy. The experimental datapoints are in this optimal region and in good agreement with the numerical simulations.

These findings were used in the experimental work described throughout this article, as the actuation voltage and *Δt* were systematically varied to achieve average step sizes spanning the range explored in simulation prior to each experiment. Guided by this characterization, all subsequent experiments were conducted using parameters that yield an average step size near the optimal minimum in the number of steps per unit distance, thereby balancing speed, accuracy, and robustness of single-cell control. The experimentally measured data, shown as red stars in Figure 2C and 2D, closely follows the simulated trends for both performance metrics, demonstrating good agreement between simulations and experiment. To directly compare experiments with numerical, *A*_*error*_ was normalized with respect to the target path length.

Substantial variability in step size across acoustic modes was observed in experimental results, reflecting the strongly mode-dependent gradients of the acoustic potential. This variability suggests that future implementations could benefit from adaptive normalization strategies, such as mode-specific modulation of actuation voltage or actuation duration *Δt*, to further improve convergence speed and trajectory fidelity.^[34]^

### 2.3. Single cell control

We experimentally demonstrate reliable two-dimensional manipulation of individual cells across multiple live cell types, including DU-145 prostate cancer cells, Jurkat T lymphocytes, and K-562 chronic myelogenous leukemia cells. These cell lines span a range of sizes, morphologies, and mechanical properties, providing a stringent test of the generality and robustness of the proposed control framework.^[41,42]^

**Figure 3**A shows snapshots of the trajectory of a single live DU-145 cell completing a task to show full 2D controllability. The target path of the trajectory was 1 mm in total, and tasks completed after 32 min. A video of the experiment can be found in Supplement B. The completed trajectory of two independent repeats performed on different experimental days and in distinct regions of the microfluidic chamber are shown in Figure 3B. They took 15 and 30 min, respectively. All experiments were conducted using identical control parameters. Equivalent experiments were conducted with Jurkat and K-562; representative trajectories for each cell type are shown in Figure 3C.

**Figure 3.**
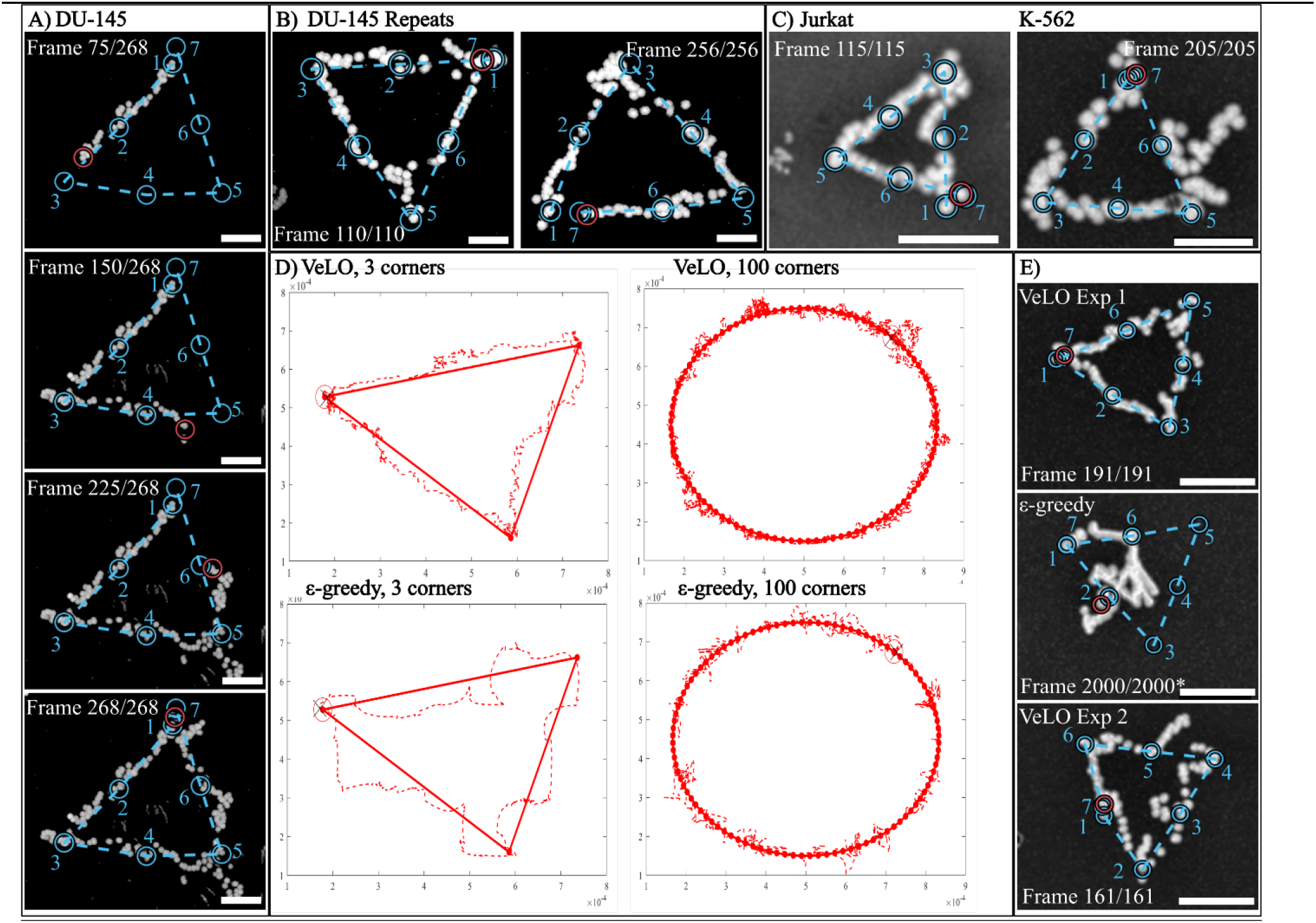
2D control of single cells. (**A**) Snapshots of the trajectory of a DU-145 cell. (**B**) Independent repetitions with DU-145 cells. (**C**) 2D control of Jurkat and K-562 cells. (**D**) Simulation comparing the VeLO algorithm with the ε-greedy algorithm. (**E**) A Jurkat cell completed a triangle with the VeLO algorithm (Exp 1) using 955 steps (191 control iterations), then attempted to repeat the same triangle with the ε-greedy algorithm. The ε-greedy experiment was aborted after 2000 steps, and the VeLO algorithm completed the triangle with the same cell again (Exp 2) using 805 steps (161 control iterations) to proof that the cell was not sticking. Note that VeLO takes 5 actuation steps for each iteration, whereas ε-greedy only takes 1 step each iteration. Numbered blue circles indicate target positions. Red circle indicates cell position. Dashed blue lines are the shortest path. Scale bars are 100 µm.

To quantitatively assess the performance of the proposed Vector-based Local Optimization (VeLO) algorithm, we performed a direct experimental comparison with an ε-greedy exploration strategy. The ε-greedy algorithm has previously been used to manipulate single 70 µm polystyrene particles using frequency-modulated standing acoustic waves.^[35–37]^

The algorithms were compared numerically by tasking them to follow paths defined by 3, 8, 20, and 100 intermediate target points. Figure 3D shows the trajectories for the simplest (3 targets) and most demanding (100 targets) cases. The complete data set can be found in Supplement C. As target point resolution increased from 3 to 100 target points, the normalized number of steps increased from 1170 to 4266 steps/mm for VeLO and from 1111 to 3476 steps/mm for ε-greedy. Over the same range, the normalized *A*_*error*_ increased modestly from 0.0207 to 0.0222 mm^2^ for VeLO, while decreasing from 0.0361 to 0.0271 mm^2^ for ε-greedy.

Figure 3E) presents an experimental comparison between algorithms performed on a single Jurkat cell. In the first phase, the cell was guided along a predefined trajectory using the VeLO algorithm, completing the task in 955 actuation steps over 191 control iterations. The experiment was then repeated using the ε-greedy algorithm. As evident from the image sequence, the cell failed to progress beyond the first of six intermediate target positions even after 2000 control iterations. To show that the cell had not adhered to the chamber surface, the experiment was again repeated using the VeLO algorithm, which completed the trajectory in 805 steps over 161 control iterations.

These results show that both algorithms perform comparably in terms of speed for low-resolution tasks with few target points (left plots in Figure 3D), with the number of steps differing 5% in favor of VeLO. However, VeLO consistently achieves higher accuracy, with a 43% smaller error area. For an increased number of intermediate targets (right plots in Figure 3D), ε-greedy begins outperforming VeLO in terms of number of steps (19% faster) and improves 25% in terms of accuracy compared to only using few target points. VeLO’s accuracy remains comparatively constant when increasing the number of target points, with an improvement of (7%), but remains 18% accurate than ε-greedy even for the simulations using large number of target points.

The discrepancy between algorithm performance observed experimentally, with ε-greedy frequently failing, indicates that VeLO is more robust to the imperfections and measurement noise that are inherent to real systems. This robustness comes from building a model based on physical phenomena such as the displacement matrix. Additionally, VeLO actively penalizes actuation modes that vary a lot, which helps avoid modes that are in a local minimum at the current cell position. As experiments were performed at frequencies matching higher order actuation fields, the distances between minima are significantly shorter than in the simulations, which utilize the lowest order actuation frequencies.

### 2.4. Independent control of multiple cells

We next demonstrate the simultaneous and independent control of multiple cells, highlighting the scalability of the VeLO algorithm beyond single-cell manipulation. In these experiments, two DU-145 prostate cancer cells were controlled concurrently within the same microfluidic chamber and guided toward a common final target position located between their initial locations. Snapshots of the trajectory of one such experiment are shown in **Figure 4**A). The experiment required 348 control iterations, corresponding to 49 minutes. Videos can be found in Supplement B. Repetitions of the experiment can be found in Supplement C.

**Figure 4.**
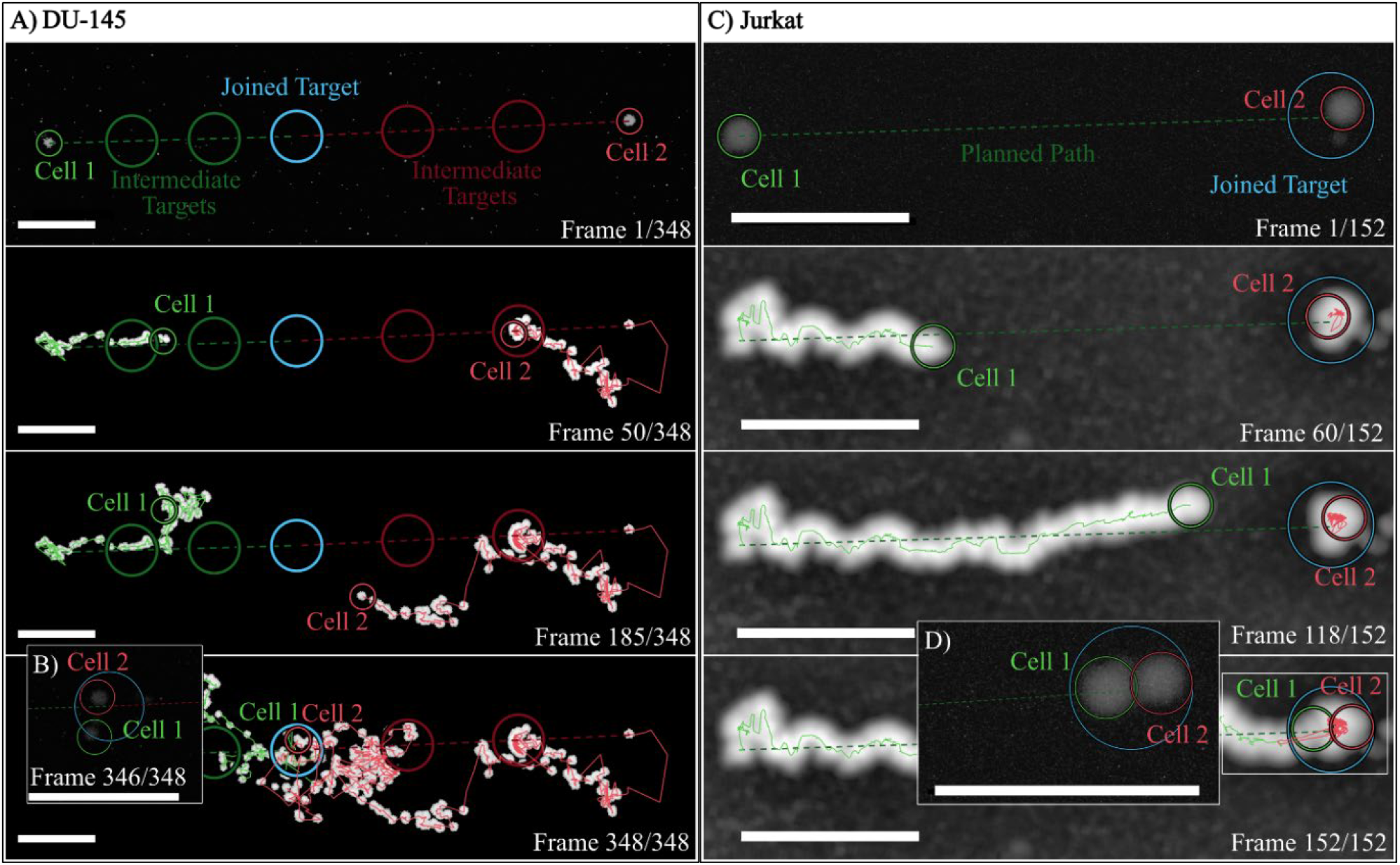
Independent, simultaneous control of two cells. (A) Two DU-145 cells (green/red small circles) past intermediate target position (large green/red) circles) towards a joint target (blue). (**B**) At frame 346, the machine vision algorithm failed to distinguish between the two cells due to proximity. The inset frame shows both cells inside the target area. (**C**) Two Jurkat cells, one (red) tasked to stay within the joined target (blue) position while the other (green) was brought towards it. Dashed lines are the shortest path. (**D**) Final frame of the experiment in C. Both cells are within the target area and in direct contact with each other. Repeats with different cell lines are shown in Supplements C. Scale bar is 100 µm.

In an additional set of experiments, one cell was tasked to remain within the target area while a second cell was actively guided towards it. This task is particularly relevant for applications involving controlled cell–cell interactions, such as immune synapse formation or signaling studies.^[43,44]^ Snapshots of one such experiment, using Jurkat T lymphocytes, are shown in Figure 4C). Additional experiments with DU-145 and K-562 leukemia cells can be found in Supplement C. This demonstrates that VeLO can selectively manipulate individual cells within a shared acoustic field and complete different tasks for each cell. Furthermore, as can be seen in Figure 4D), this method enables bringing two cells into direct contact with each other. To our knowledge, the controlled merging or enforced contact of individual cells using multimodal acoustofluidic feedback has not been previously demonstrated.

## 3. Discussion

In this work, we introduce a closed loop acoustofluidic control framework for programmable manipulation of individual living cells, based on a locally optimizing feedback algorithm termed VeLO. By dynamically modulating standing acoustic wave fields and learning system dynamics online, VeLO enables precise two-dimensional control of single cells without physical contact, labels, or prior system calibration. To our knowledge, this represents the first demonstration of feedback-controlled, single-cell–level manipulation using standing-wave acoustofluidics. We show that VeLO generalizes across different cell types, including adherent-derived cancer cells and suspension lymphocytes, despite differences in size, morphology, and mechanical properties. The numerical simulations accurately predicted optimal operating conditions, showing that a tradeoff exists between the average step size and the accuracy of control, which will be an important tool in optimizing hyperparameters for various future applications.

Furthermore, the potential for multicell handling, including controlled cell–cell contact, is explored in this work by independently controlling two cells simultaneously, bringing both to a common target position or keeping one at its starting position while bringing the other cell to it. The presented technology offers a versatile platform for biological investigations requiring precise, non-invasive control of individual cells. Applications range from controlled cell-cell interactions, deterministic single-cell seeding for tissue engineering and organoid formation, and selective extraction or relocation of rare cells from heterogeneous populations. Since acoustic manipulation is inherently gentle, label-free, and compatible with standard microfluidic environments, the approach is well suited for integration into existing lab-on-a-chip workflows.

## 4. Materials and methods

### 4.1. Platform and device

The platform consists of a function generator (Tektronix AFG 3022B), which is connected to the microfluidic device through an in-house built amplifier.

A microscope (Leica DM2500M) and a camera (IDS U3-3890CP Rev2.2) are used for visual feedback to the control algorithm. An X-cite series 120 Q light source is used with a standard fluorescent green filter cube (excitation peak at 490 nm; emission peak at 525 nm) provided fluorescent light. The camera and function generator are connected to a computer with USB-A cables.

The microfluidic device is an anodically bonded glass-silicon and was manufactured at the Danish Technical University, DTU. The width and length of the device is 8.8 x 20.25 mm. The glass layer is 500 µm thick, with 150 µm wide holes for fluid connections. The silicon layer is 300 µm thick, and the fluidic chamber is 150 µm deep and dry etched into the silicon layer. The chamber’s bottom shape is an irregular hexagon with a surface area of 13.6 mm^2^. 2 channels with a width of 200 µm connect the chamber to the inlet holes.

The piezoelectric transducer (Pz26; Meggit Ferroperm Piezoceramics) is 4.9 x 12.9 x 0.4 mm and was glued (Loctite Super Glue Brush on) to a bottom corner of the device. Two silicon tubes (outer diameter 3 mm, inner diameter 1 mm) are glued to the glass layer at the inlet holes and allowed to fill the device up with help of 1ml syringes.

### 4.2. Cell culturing and preparation

Human prostate cancer cell DU-145, Jurkat T lymphocytes, and K-562 myelogenous leukemia cell lines were acquired from American Type Culture Collection and grown according to the supplier’s recommendations.

Prior to experiments, the device was coated with 1% Pluronic F-127 for 30 minutes to reduce cell adhesion. Cells were stained by adding 4 µL calcein AM to a suspension of 10^6^ cells in 1 mL cell buffer for 20 minutes on ice. Excessive calcein was removed by adding 9 mL cell buffer, centrifuging for 5 min at 400 rpm, removal of the buffer and resuspension with fresh cell buffer. Subsequentially, the cell suspension was diluted until a suitable cell concentration was found. Cells were introduced to the manipulation device through the device inlet with a syringe.

### 4.3. Experimental procedure

The cell suspension was perfused into the microfluidic device through the inlet hole, taken care to not create bubbles inside of the microfluidic chamber. After controlling the mobility of the cells, and selecting suitable input parameters (voltage, actuation time, frequency range and number of modes), the target cell/s was selected by drawing a rectangle around it with help of a custom Matlab GUI. The target trajectory was created by manually selecting target positions in the GUI. Once cells and targets were selected, the experiment was initiated, running autonomously controlled by the chosen control algorithm until a termination condition was fulfilled.

### 4.4. Machine vision

The machine vision in the setup uses a local cross correlation paired with a nearest neighbor approach. The cell is initially selected by drawing a rectangle around it. The machine vision identifies the cell inside of the rectangle by finding the largest cluster of brighter pixels and saves its approximate size and shape as a template. For each tracking frame, the machine vision algorithm searches for objects matching the template within a defined region around the previous cell position. Should multiple cells be identified, the cell closest to the previous position is tracked.

### 4.5. Numerical results

The numerical simulations were set up to model the movement of a particle in the lower left quarter of a rectangular chamber. During simulations, the particles’ initial and target positions were manually defined before the experiment. The control logic was the same as used in the experimental platform. The particle displacement was derived by first generating the potential field in the chamber corresponding to the selected actuation frequency and calculating the force vector on each cell. The vector multiplied with *Δt* gave the cell displacement. All numerical data is available in Supplement E.

## Funding and Acknowledgements

Funding: T. B. is supported by the Swedish Research Council (No. 2022-04041), the Crafoordska Stiftelsen (No. 20240891), and the SONOCRAFT project funded by the European Innovation and Research Council (GA: 101187842). T.B. and A.E. acknowledge support from the Life Science Microfluidics Infrastructure at Lund University and the Engineering Health profile area. A.E. is supported by the Swedish Research Council (No. 2019-00795 & 2022-04041), Prostatacancerförbundet and Fru Berta Kamprad Stiftelse (No. FBKS-2022-34-(426)). We would like to acknowledge Prof. Thomas Laurell and our co-workers in the Acoustofluidics Group at the Department of Biomedical Engineering for their input and support.

## Data Availability Statement

All data needed to evaluate the conclusions in the paper are present in the paper and/or the Supplementary Materials. The MATLAB framework is available upon reasonable request to the corresponding author.

