## Supplements for "Programmable acoustic single cell manipulation with model-free machine learning"

**Supplement A: COMSOL simulation**

The acoustic pressure field inside the microfluidic chamber was modeled in COMSOL Multiphysics (v6.3) using the *Pressure Acoustics, Frequency Domain* interface. The simulated geometry replicated the chamber dimensions of the experimental device, see the technical drawing in Supplement B for dimensions. All solid boundaries were modeled as acoustically rigid (sound hard boundary condition), corresponding to high acoustic impedance mismatch between the fluid (PBS) and the surrounding solid materials.

The fluid domain was assigned the material properties of phosphate-buffered saline (PBS), with density $\rho_0$ = 1004 kg/m$^3$ and speed of sound $c_0$ = 1508 m/s. The particles were modeled using parameters corresponding to DU145 cells, assuming spherical geometry with radius $a$ = 7.5 µm, density $\rho_c$ = 1062 kg/m$^3$, and compressibility $\kappa_c$ = 384 T/Pa.[39]

An eigenfrequency study was performed using the ARPACK solver to compute the first 200 acoustic eigenmodes of the fluid domain. The computed eigenfrequencies spanned the range from 0.18 MHz to 4.07 MHz. A shift-invert method was employed to ensure convergence in the frequency region of interest. The eigenmodes were normalized to unit maximum pressure amplitude.

The acoustic radiation potential (Gor'kov potential) was calculated for each eigenmode using Equation. 2. The acoustic velocity field was obtained from the pressure gradient via

$$\boldsymbol{v} = -\frac{1}{i\omega\rho_0}\boldsymbol{\nabla}p. \tag{S1}$$

The mesh was refined to a minimum element size of 6 µm and a maximum element size of 0.12 µm, resulting in approximately 22915 degrees of freedom.

The model assumes linear acoustics, neglects viscous boundary layers and acoustic streaming effects, and is valid under the small particle approximation (a ≪ λ).

The COMSOL model file is available from the corresponding author upon reasonable request.

**Supplement B: Videos**

Power point “Video Supplements” with video files:
“01_DU145_LTH.mp4”
“02_DU145_Triangle.mp4”
“03_DU145_2cells.mp4”
“04_Jurkat_2Cells.mp4”

Are available upon reasonable request.

**Supplement C: Experimental Data**

**Table S1:** Overview of all experimental data. The colored lines in the trajectory image represent different frequencies that were applied, showing how the algorithm switches between them.

| Experiment | Parameters | Data Results | Trajectory Image |
|---|---|---|---|
| Du-145,<br>VeLO,<br>Triangle | $V_{in}$: 14 V<br>Δt: 0.4 s<br>StepSize: 3.55 μm<br>Modes: 30<br>Range: 1.89-2.85 MHz<br>λ: 0.0005<br>$\gamma$: 0.8 | Iterations: 110<br>Steps/mm: 530<br>$A_{error}$: 11770 μm$^2$<br>Time: 15 min | 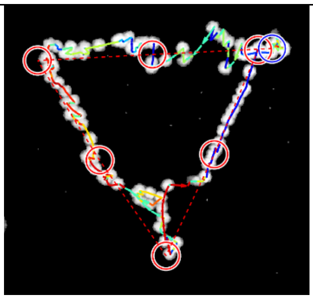 |
| Du-145,<br>VeLO,<br>Triangle | $V_{in}$: 14 V<br>Δt: 0.5 s<br>StepSize: 1.90 μm<br>Modes: 30<br>Range:1.89-2.85 MHz<br>λ: 0.0005<br>$\gamma$: 0.8 | Iterations: 617<br>Steps/mm: 2820<br>$A_{error}$: 57340 μm$^2$<br>Time: 72 min | 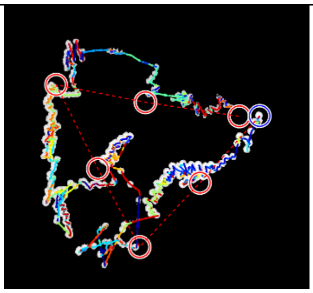 |
| Du-145,<br>VeLO,<br>Triangle | $V_{in}$: 14 V<br>Δt: 0.4 s<br>StepSize: 2.63 μm<br>Modes: 30<br>Range:1.89-2.85 MHz<br>λ: 0.0005<br>$\gamma$: 0.8 | Iterations: 268<br>Steps/mm: 1250<br>$A_{error}$: 17970 μm$^2$<br>Time: 32 min | 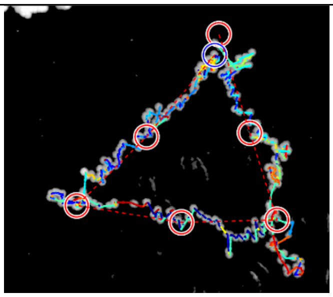 |
| Du-145,<br>VeLO,<br>Triangle | $V_{in}$: 14 V<br>Δt: 0.4 s<br>StepSize: 3.62 μm<br>Modes: 30<br>Range:1.89-2.85 MHz | Iterations: 399<br>Steps/mm: 1792<br>$A_{error}$: 128720 μm$^2$<br>Time: 47 min | 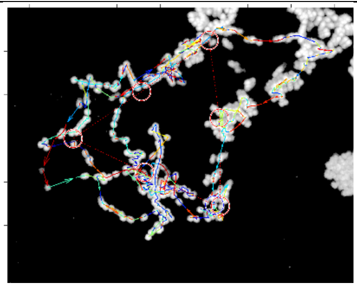 |

| | | | |
|---|---|---|---|
| | λ: 0.0005<br>$\gamma$: 0.8 | | |
| Du-145,<br>VeLO,<br>Triangle | $V_{in}$: 14 V<br>Δt: 0.4 s<br>StepSize: 2.38 µm<br>Modes: 30<br>Range:1.89-2.85 MHz<br>λ: 0.0005<br>$\gamma$: 0.8 | Iterations: 457<br>Steps/mm: 2098<br>$A_{error}$: 34100 µm$^2$<br>Time: 54 min | 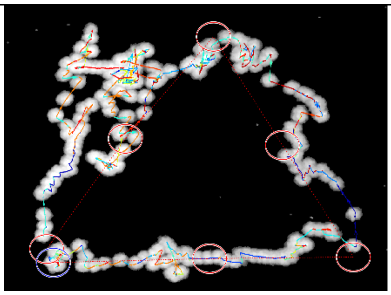 |
| Du-145,<br>VeLO,<br>Triangle | $V_{in}$: 14 V<br>Δt: 0.4 s<br>StepSize: 2.38 µm<br>Modes: 30<br>Range:1.89-2.85 MHz<br>λ: 0.0005<br>$\gamma$: 0.8 | Iterations: 256<br>Steps/mm: 1197<br>$A_{error}$: 16800 µm$^2$<br>Time: 30 min | 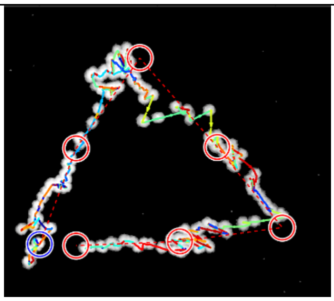 |
| Du-145,<br>VeLO,<br>Two-cells | $V_{in}$: 23 V<br>Δt: 0.6 s<br>StepSize: µm<br>Modes: 40<br>Range: 2.0 – 3.2 MHz<br>λ: 0.0005<br>$\gamma$: 0.8 | Iterations: 276<br>Steps/mm:<br>$A_{error}$: µm$^2$<br>Time: 46 min | 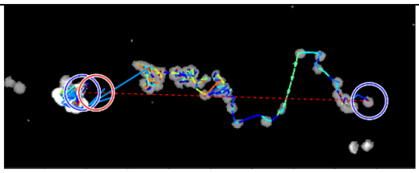 |
| Du-145,<br>VeLO,<br>Two-cells | $V_{in}$: 20 V<br>Δt: 0.7 s<br>StepSize: µm<br>Modes: 40<br>Range: 2.0 – 3.2 MHz<br>λ: 0.0005<br>$\gamma$: 0.8 | Iterations: 86<br>Steps/mm:<br>$A_{error}$: µm$^2$<br>Time: 14 min | 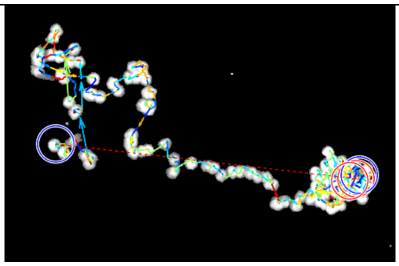 |

| | | | |
|---|---|---|---|
| Du-145,<br>VeLO,<br>Two-cells | $V_{in}$: 18 V<br>Δt: 0.3 s<br>StepSize: µm<br>Modes: 40<br>Range: 1.5 – 2.8 MHz<br>λ: 0.0005<br>$\gamma$: 0.2 | Iterations: 362<br>Steps/mm:<br>$A_{error}$: µm$^2$<br>Time: 49 min | 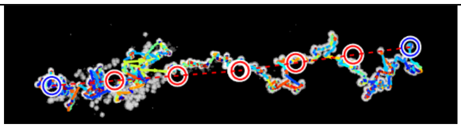 |
| Du-145,<br>VeLO,<br>Two-cells | $V_{in}$: 18 V<br>Δt: 0.3 s<br>StepSize: µm<br>Modes: 40<br>Range: 1.5 – 2.8 MHz<br>λ: 0.0005<br>$\gamma$: 0.2 | Iterations: 349<br>Steps/mm:<br>$A_{error}$: µm$^2$<br>Time: 53 min | 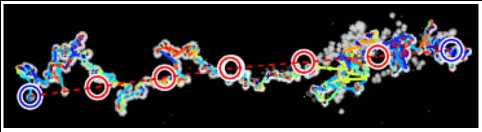 |
| Du-145,<br>VeLO,<br>Two-cells | $V_{in}$: 18 V<br>Δt: 0.3 s<br>StepSize: µm<br>Modes: 40<br>Range: 1.5 – 2.8 MHz<br>λ: 0.0005<br>$\gamma$: 0.2 | Iterations: 141<br>Steps/mm:<br>$A_{error}$: µm$^2$<br>Time: 22 min | 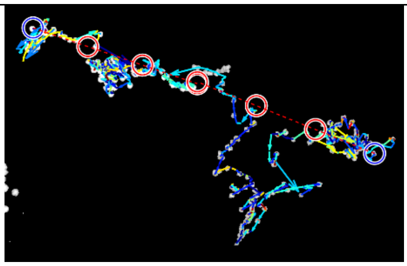 |
| Du-145,<br>VeLO,<br>Letter L | $V_{in}$: 20 V<br>Δt: 0.3 s<br>StepSize: 1.90 µm<br>Modes: 40<br>Range: 1.5 – 2.8 MHz<br>λ: 0.0005<br>$\gamma$: 0.2 | Iterations: 163<br>Steps/mm: 1524<br>$A_{error}$: 23060 µm$^2$<br>Time: 31 min | 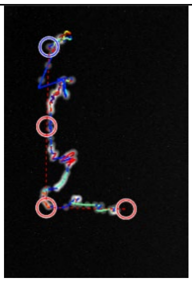 |

| | | | |
|---|---|---|---|
| Du-145,<br>VeLO,<br>Letter T | $V_{in}$: 20 V<br>Δt: 0.3 s<br>StepSize: 1.69 μm<br>Modes: 40<br>Range: 1.5 – 2.8 MHz<br>λ: 0.0005<br>$\gamma$: 0.2 | Iterations: 258<br>Steps/mm: 1158<br>$A_{error}$: 705160 μm$^2$<br>Time: 44 min | 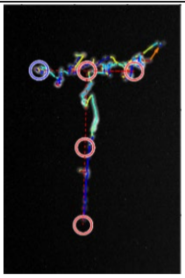 |
| Du-145,<br>VeLO,<br>Letter H | $V_{in}$: 20 V<br>Δt: 0.3 s<br>StepSize: 1.73 μm<br>Modes: 40<br>Range: 1.5 – 2.8 MHz<br>λ: 0.0005<br>$\gamma$: 0.2 | Iterations: 446<br>Steps/mm: 1725<br>$A_{error}$: 573537 μm$^2$<br>Time: 76 min | 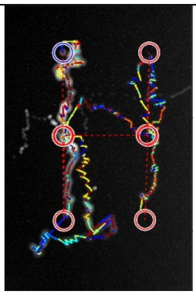 |
| Jurkat<br>VeLO<br>Triangle | $V_{in}$: 14 V<br>Δt: 0.3 s<br>StepSize: 1.37 μm<br>Modes: 40<br>Range: 1.5 – 2.8 MHz<br>λ: 0.0005<br>$\gamma$: 0.2 | Iterations: 161<br>Steps/mm: 1769<br>$A_{error}$: 6401 μm$^2$<br>Time: 25 min | 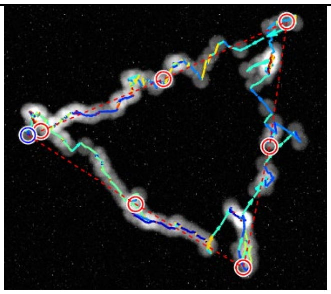 |
| Jurkat<br>VeLO<br>Triangle | $V_{in}$: 20 V<br>Δt: 0.5 s<br>StepSize: 2.10 μm<br>Modes: 30<br>Range: 1.3 – 2.8 MHz<br>λ: 0.0005<br>$\gamma$: 0.8 | Iterations: 109<br>Steps/mm: 1389<br>$A_{error}$: 5897 μm$^2$<br>Time: 14 min | 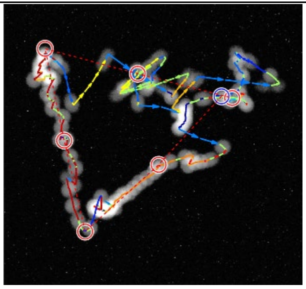 |

| | | | |
|---|---|---|---|
| Jurkat<br>VeLO<br>Triangle | $V_{in}$: 20 V<br>Δt: 0.3 s<br>StepSize: 3.13 μm<br>Modes: 30<br>Range:  1.3 – 2.8 MHz<br>λ: 0.0005<br>$\gamma$: 0.8 | Iterations: 464<br>Steps/mm: 5616<br>$A_{error}$: 6400 μm$^2$<br>Time: 55 min | 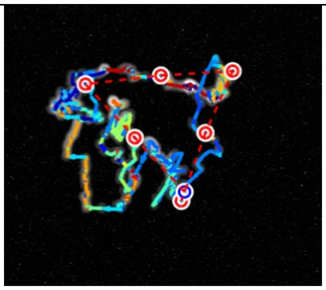 |
| Jurkat<br>VeLO<br>Triangle | $V_{in}$: 20 V<br>Δt: 0.5 s<br>StepSize: 1.09 μm<br>Modes: 30<br>Range:  1.3 – 2.8 MHz<br>λ: 0.0005<br>$\gamma$: 0.8 | Iterations: 105<br>Steps/mm: 1339<br>$A_{error}$: 7684 μm$^2$<br>Time: 14 min | 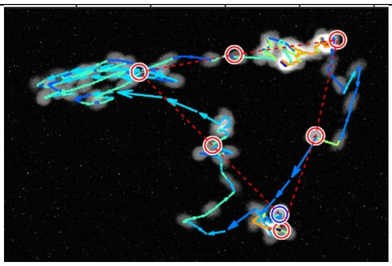 |
| Jurkat<br>VeLO<br>Triangle | $V_{in}$: 14 V<br>Δt: 0.3 s<br>StepSize: 1.82 μm<br>Modes: 40<br>Range:  1.5 – 2.8 MHz<br>λ: 0.0005<br>$\gamma$: 0.2 | Iterations: 191<br>Steps/mm: 1527<br>$A_{error}$: 6619 μm$^2$<br>Time: 26 min | 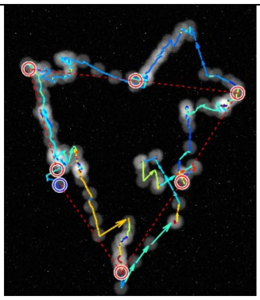 |
| Jurkat<br>ε-greedy Triangle | $V_{in}$: 14 V<br>Δt: 0.3 s<br>StepSize: 0.69 μm<br>Modes: 40<br>Range:  1.5 – 2.8 MHz<br>ε: 0.1<br>$\theta$= 0.999 | Iterations: 2000+<br>Steps/mm: 21786<br>$A_{error}$: 5868 μm$^2$<br>Time: 84 min<br>ABORTED after 1/6$^{th}$ of trajectory | 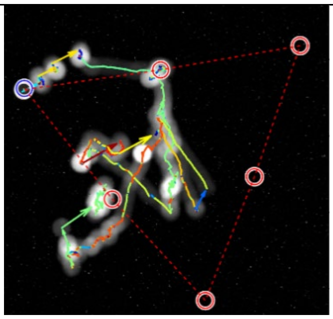 |

| | | | |
|---|---|---|---|
| Jurkat<br>VeLO<br>2 cells | $V_{in}$: 10 V<br>Δt: 0.5 s<br>StepSize: μm<br>Modes: 30<br>Range: 1.3 – 2.8 MHz<br>λ: 0.0005<br>$\gamma$: 0.8 | Iterations: 152<br>Steps/mm:<br>$A_{error}$: μm$^2$<br>Time: 29 min | 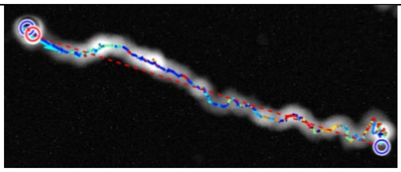 |
| K-562<br>VeLO<br>Triangle | $V_{in}$: 10 V<br>Δt: 0.3 s<br>StepSize: 0.89 μm<br>Modes: 30<br>Range: 1.8 – 2.8 MHz<br>λ: 0.0005<br>$\gamma$: 0.8 | Iterations: 403<br>Steps/mm: 4799<br>$A_{error}$: 4095 μm$^2$<br>Time: 54 min | 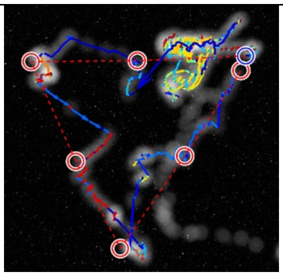 |
| K-562<br>VeLO<br>Triangle | $V_{in}$: 10 V<br>Δt: 0.3 s<br>StepSize: 1.74 μm<br>Modes: 30<br>Range: 1.8 – 2.8 MHz<br>λ: 0.0005<br>$\gamma$: 0.8 | Iterations: 136<br>Steps/mm: 1666<br>$A_{error}$: 7396 μm$^2$<br>Time: 44 min | 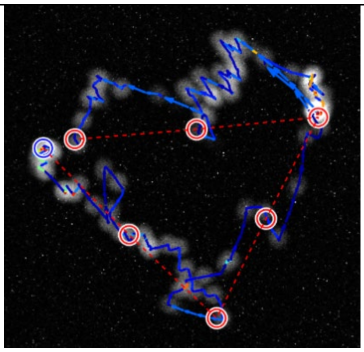 |
| K-562<br>VeLO<br>Triangle | $V_{in}$: 18 V<br>Δt: 0.3 s<br>StepSize: 2.35 μm<br>Modes: 30<br>Range: 1.8 – 2.8 MHz<br>λ: 0.0005<br>$\gamma$: 0.2 | Iterations: 205<br>Steps/mm: 1882<br>$A_{error}$: 15009 μm$^2$<br>Time: 38 min | 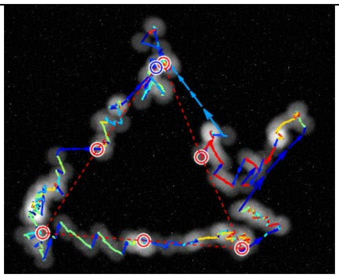 |

| | | | |
|---|---|---|---|
| K-562<br><br>ε-greedy Triangle | $V_{in}$: 18 V<br><br>Δt: 0.3 s<br><br>StepSize: μm<br><br>Modes: 30<br><br>Range: 1.85 – 2.8 MHz<br><br>ε: 0.1<br><br>$\theta$= 0.9 | Iterations: 1410<br><br>Steps/mm:<br><br>$A_{error}$: μm$^2$<br><br>Time: 60 min<br><br>ABORTED, stuck | 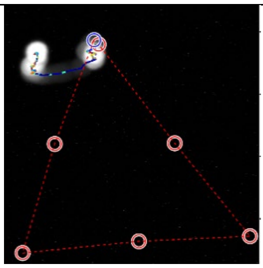 |
| K-562<br><br>ε-greedy Triangle | $V_{in}$: 18 V<br><br>Δt: 0.3 s<br><br>StepSize: μm<br><br>Modes: 30<br><br>Range: 1.85 – 2.8 MHz<br><br>ε: 0.1<br><br>$\theta$= 0.9 | Iterations: 1159<br><br>Steps/mm:<br><br>$A_{error}$: μm$^2$<br><br>Time: 54 min<br><br>ABORTED, stuck | 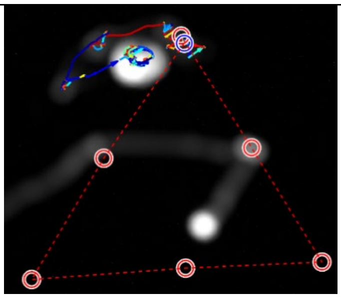 |
| K-562<br><br>ε-greedy Triangle | $V_{in}$: 10 V<br><br>Δt: 0.3 s<br><br>StepSize: μm<br><br>Modes: 30<br><br>Range: 1.85 – 2.8 MHz<br><br>ε: 0.1<br><br>$\theta$= 0.999 | Iterations: 2000+<br><br>Steps/mm:<br><br>$A_{error}$: μm$^2$<br><br>Time: 62 min<br><br>ABORTED | 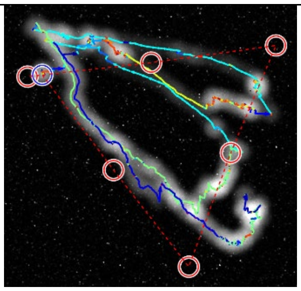 |
| K-562<br><br>VeLO<br><br>2 cells | $V_{in}$: 10 V<br><br>Δt: 0.4 s<br><br>StepSize: μm<br><br>Modes: 30<br><br>Range: 1.8 – 2.8 MHz<br><br>λ: 0.0005<br><br>$\gamma$: 0.8 | Iterations: 54<br><br>Steps/mm:<br><br>$A_{error}$: μm$^2$<br><br>Time: 10 min | 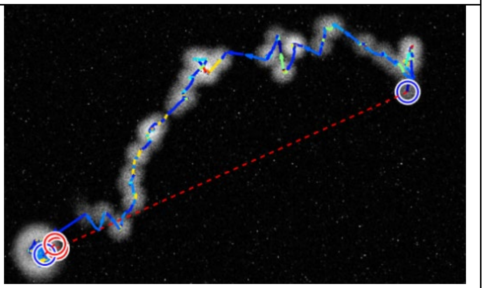 |

**Supplement D: Numerical model**

All simulations were performed in MATLAB R2024A within a two-dimensional rectangular domain of dimensions L = 1.6mm and W = 1.9mm. The model assumes planar standing waves and neglects out-of-plane variations. Boundary conditions were imposed as acoustically rigid walls with pressure $p_x(0, L) = 0$, and $p_y(0, W) = 0$, where x and y denote the x and y derivative, respectively.

The acoustic pressure field corresponding to mode indices $(n, m)$ was defined analytically as

$$p_{n,m}(x, y) = p_0 \cos\left(\frac{n\pi x}{L}\right) \cos\left(\frac{m\pi y}{\mathrm{W}}\right), \tag{S2}$$

where $n, m \in \{1, \ldots, 6\}$ and $p_0$ is the pressure amplitude. The action space therefore comprised 36 distinct spatial mode patterns. The associated first-order acoustic velocity field was obtained from the linearized momentum equation, Eq. S2.

The time-averaged acoustic radiation force was derived from the Gorkov potential, Eq. 2. Fluid properties were chosen to approximate PBS: $c_0 = 1508$ m/s, $\rho_0 = 1004$kg/m$^3$, $\kappa_0 = 438$ T/Pa, and $\eta_0 = 0.001$Pa·s. The particle was modeled as a spherical DU-145 cell of radius $a = 7.5$ µm with acoustic contrast factors $f_1 = 0.12$ and $f_2 = 0.038$. Values obtained from literature[39].

Particle motion was governed by balancing the acoustic radiation force and Stoke's drag force:

$$\boldsymbol{v} = \frac{\boldsymbol{F}_{rad}}{6\pi\eta_0 a}. \tag{S3}$$

Inertia, brownian motion, acoustic streaming, secondary radiation forces, wall corrections, and 3D effects were neglected.

Trajectory integration was performed using MATLAB's ode45 solver. The physical integration time step was constrained to $\Delta t_{dyn} = 0.01$s. The control algorithm operated at $\Delta t_{control} = 0.5$s, such that each control step consisted of 50 dynamic updates.

At each control iteration $k$, given particle position $\mathrm{x}_k$ and target position $\mathrm{x}_t$, a mode or mode mixture was selected using ε-greedy or VeLO policy. The acoustic field was applied for $\Delta t_{control}$ for each mode (or each mode in the mode mixture), the particle trajectory was integrated, and the updated position was returned as feedback. The reward function for ε-greedy was defined as $r = \| x_{particle,k-1} - x_{target} \|_2 - \| x_{particle,k} - x_{target} \|_2$, which updated the weight $\mu$ according to: $\mu(n, m) = \mu(n, m) + \left(r - \frac{\mu(n,m)}{w(n,m)}\right)$, with $w$ being a normalization term.

For the numerical characterization of the VeLO policy, the pressure amplitude $p_0$ was incrementally varied over the range [27000‚ 205000]in steps of $\Delta p$ =[5400]. For each amplitude, the number of control steps required to complete a 1 mm triangular trajectory was recorded. The sweep was terminated when the required steps exceeded 7200 (corresponding to 1 hour of actuation). Performance metrics included mean step size, number of steps for completion, error area $A_{error}$.

A target was considered reached when

$$\| \mathrm{x}_{particle} - \mathrm{x}_{target} \|_2 < a. \tag{S4}$$

All MATLAB code is available from the corresponding author upon reasonable request.

**Supplement E: Numerical Comparison of VeLO and $\epsilon$-greedy data**

Table S2: Numerical Results of the algorithm comparison.

| **Algorithm** | **Target Points** | **Average Displacement (µm)** | **Steps/mm** | **$A_{error}$/mm ($10^{-8}$ $m^2$)** |
|---|---|---|---|---|
| VeLO | 3 | 3.46 | 1170 | 2.07 |
| VeLO | 8 | 3.35 | 1253 | 1.68 |
| VeLO | 20 | 3.22 | 1844 | 1.18 |
| VeLO | 100 | 3.05 | 4266 | 2.22 |
| $\epsilon$-greedy | 3 | 2.00 | 1111 | 3.61 |
| $\epsilon$-greedy | 8 | 2.12 | 1190 | 2.94 |
| $\epsilon$-greedy | 20 | 2.02 | 1549 | 1.79 |
| $\epsilon$-greedy | 100 | 1.79 | 3476 | 2.71 |
